# Proteomic profiling of skeletal muscle ribosomes from higher versus lower responders to 10 weeks of resistance training

**DOI:** 10.1101/2025.11.04.686603

**Authors:** J. Max Michel, Samuel C. Norton, João G. A. Bergamasco, Maíra C. Scarpelli, Talisson S. Chaves, Deivid G. da Silva, Diego Bittencourt, Chidozie G. Ugochukwu, Melissa D. Boersma, C. Brooks Mobley, Joshua S. Godwin, Gustavo A. Nader, Cleiton A. Libardi, Michael D. Roberts

**Author notes:** address correspondence to Michael D. Roberts, PhD, Endowed Alumni Professor, Director, Nutrabolt Applied and Molecular Physiology Laboratory, School of Kinesiology, Auburn University.

## Abstract

Ribosome biogenesis is a key driver of resistance training (RT)-induced skeletal muscle hypertrophy in humans. However, high resolution insight into RT-induced compositional alterations in ribosomes remain unexplored. Therefore, the purpose of this study was twofold: 1) develop a protocol sufficient to enrich and examine the ribosomal proteome from a small amount of human muscle tissue, and 2) determine if ribosomal protein composition differs between higher and lower responders to RT. Fourteen participants completed 10 weeks of RT (23 sessions) and were stratified into higher (n=7) and lower (n=7) responders based on changes in vastus lateralis muscle cross-sectional area (VL mCSA) and mean myofiber cross-sectional area (fCSA) from baseline (PRE) to after the 23^rd^ RT session (POST). Participants then performed a twenty fourth RT session and biopsies were collected 24 hours post-RT (POST-24h) to examine acute bout (POST to POST-24h) ribosomal proteome alterations to RT. Ribosome enrichment and analysis included ultracentrifugation (100,000 *g*, 3h, 2°C) through 20% sucrose gradients followed by shotgun proteomics to quantify ribosomal protein composition. Our protocol produced exceptional ribosome enrichment, with 74 distinct ribosomal proteins being detected (92% coverage of the putative 80 cytosolic ribosomal proteins) as well as ∼164-fold and ∼71-fold increases in large (RPL) and small (RPS) protein subunit abundances, respectively, compared to conventional tissue homogenization. Despite robust phenotypic differences in hypertrophy between responder groups to the 10-week RT protocol (mCSA: +31% versus +3%; fCSA: +29% versus -2%, P<0.05 for both outcomes), ribosomal protein composition was not significantly different between groups (higher vs. lower responders), nor was there a significant time (POST to POST-24h) or a group*time interaction (P>0.028 for all comparisons). In conclusion, we present a highly effective ribosome enrichment protocol requiring minimal human muscle tissue. Though preliminary analyses indicate that ribosomal protein composition was similar between hypertrophic phenotypes in response to an acute RT session after a 10-week RT intervention, future research leveraging our techniques with more sampling time points are needed to provide more definitive conclusions.

## INTRODUCTION

Skeletal muscle is a plastic tissue capable of responding to external stimuli (Fluck & Hoppeler, 2003). The hallmark adaptation of skeletal muscle to mechanical overload is the enlargement of myofibers, or hypertrophy, (Roberts *et al*., 2023). While this is a well-established phenotypic response, the degree to which individuals respond to resistance training (RT) is highly variable (Bamman *et al*., 2007; Mann *et al*., 2014). This heterogenic interindividual response (Sparks, 2017; Lavin *et al*., 2021) led to the investigation of molecular mechanisms differentiating those that demonstrate robust hypertrophy with RT (higher responders) from those exhibiting less evident hypertrophic adaptation (lower responders) to RT (Roberts *et al*., 2018a; Lixandrao *et al*., 2025; McAdam *et al*., 2025). Ribosomal adaptations have been consistently associated with muscle growth (Nader *et al*., 2005; Stec *et al*., 2016; Kim *et al*., 2019), and an increase in ribosome content (determined by total RNA assessment) occurs in parallel with RT-induced hypertrophy (Kadi *et al*., 2004; Figueiredo *et al*., 2015; Reidy *et al*., 2017; Michel *et al*., 2025). Deuterium oxide-based tracer methods have additionally been used to show that ribosome synthesis rates increase with RT in young healthy males (Brook *et al*., 2017). Furthermore, ribosome content increases have been shown to occur in higher (but not lower) hypertrophic responders to RT in both younger and older adults (Stec *et al*., 2016; Mobley *et al*., 2018).

More nuanced ribosome alterations have also been investigated in the context of muscle growth, albeit this line of research has been limited. For instance, *in vitro* analysis has shown that the preferential expression of the ribosomal protein L3-like (RPL3L) impairs myoblast fusion leading to decrements in myotube size (Chaillou *et al*., 2016). This finding suggests that individual ribosomal proteins may play key roles in the regulation of skeletal muscle size, and supports the notion that specialized ribosomes may modulate the hypertrophic response to RT. It is also notable that ribosomal heterogeneity can be achieved through post-translational modifications of ribosomal proteins (Liew & Gornall, 1973; Shin *et al*., 2009; Shirai *et al*., 2010; Matsuo *et al*., 2017; An & Harper, 2020; Montellese *et al*., 2020; Pletnev *et al*., 2022; Ni *et al*., 2023), changes in ribosomal protein composition (Chaillou *et al*., 2016; Chaillou, 2019), and/or incorporation of alternate forms of rRNA (Parks *et al*., 2018; Rothschild *et al*., 2024), ultimately effecting translational output in distinct ways.

While these mechanistic insights are promising, most studies on ribosomal function and specialization have been conducted using *in vitro* systems or rodent models (Mauro & Edelman, 2002, 2007; Chaillou *et al*., 2016; Chaillou, 2019). In human skeletal muscle, direct interrogation of ribosome protein composition remains technically challenging. Standard ribosome isolation protocols, such as polysome profiling, typically require large tissue inputs and are not well-suited for the small biopsy samples commonly obtained in human studies. Therefore, there is a growing need for a streamlined and cost-effective method for ribosome enrichment from limited human muscle tissue. Developing such a protocol would facilitate proteomic analyses of ribosomal composition and post-translational modifications in clinical and exercise-based human research. Moreover, despite such ribosomal modifications regulating protein synthesis rates and preferential mRNA translation (Shi *et al*., 2017; Riba *et al*., 2019), no studies have sought to examine if ribosomal protein profiles are associated with the response heterogeneity to RT.

Therefore, the purpose of this study was two-fold. First, we sought to develop a method to enrich ribosomes from human skeletal muscle. Next, we sought to characterize the ribosomal protein profile from a cohort of previously untrained individuals displaying higher or lower hypertrophic responsiveness to RT. We hypothesized that ribosomal proteins could be effectively enriched using relatively small amount of flash frozen human skeletal muscle. Additionally, we hypothesized that higher responders to RT would present a unique ribosomal protein signature prior to and 24 hours (24h) following an acute RT bout compared to their lower responder counterparts.

## METHODS

### Participants and ethical approval

Human tissue for this methodological study and secondary analysis was available from our previously published study by Scarpelli et al. (Scarpelli *et al*., 2024). In the original study, thirty-eight untrained individuals met inclusion criteria and agreed to donate muscle biopsies prior to the intervention (PRE), 24 hours following the first training bout (PRE-24h), following 23 training sessions over a 10-week period (POST), and 24 hours following the 24^th^ training session (POST-24h). Given limited tissue availability, only POST and POST-24h samples from 14 participants were used as described in later sections. The parent study described by Scarpelli et al. (Scarpelli *et al*., 2024) was registered as a clinical trial (Brazilian Registry of Clinical Trials -RBR – 57v9mrb) and approved by the Ethics Committee of the Federal University of São Carlos (no. 56259622.0.0000.5504) in accordance with the most recent version of the Declaration of Helsinki.

### Participant selection for responder analysis

In this study, a subset of 7 higher (sex = 4M/3F; age = 25±4 years; body mass = 73.0±14.6 kg; height = 1.7±0.1 m; BMI = 26.0±5.2 kg/m^2^) and 7 lower responders (sex = 3M/4F; age = 23±2 years; body mass = 70.5±14.4 kg; height = 1.7±0.1 m; BMI = 23.8±3.3 kg/m^2^) to the 10 weeks of RT were included. These 14 participants were identified as higher or lower responders based on a composite score encompassing both ultrasound-derived muscle cross-sectional area (mCSA) and fiber cross-sectional area (fSCA) of the vastus lateralis (VL). The composite score was calculated by multiplying the VL mCSA change score (from PRE to POST) by 2/3 and the fCSA change score (from PRE to POST) by 1/3. These composite scores were then rank ordered and the seven highest and seven lowest scores were selected to represent higher and lower responders respectively. Our rationale for this multi-factorial stratification of RT response heterogeneity was driven by our past publications in this area where: i) similar multi-factorial stratification methods were used (Roberts *et al*., 2018b; Godwin *et al*., 2023; Smith *et al*., 2023), and ii) we have demonstrated that different assessments of skeletal muscle hypertrophy (e.g., changes in VL fCSA and VL mCSA) show poor agreement with one another (Ruple *et al*., 2022), thus justifying the utilization of multiple variables when parsing out RT response heterogeneity.

### Resistance training protocol

A one repetition maximum (1RM) test was utilized to determine the load used in the first training session (80% 1RM). The RT protocol used for the samples herein was comprised of 4 sets of 9-12 repetitions of the leg extension exercise, performed unilaterally to concentric muscle failure. Ninety seconds of rest was taken between each set, and the load was adjusted such that participants reached concentric muscle failure within the target repetition range. Training sessions were carried out 2-3 times per week for 10 weeks totaling 23 unilateral RT sessions.

### Ultrasound imaging for muscle cross-sectional area

VL mCSA was assessed by an experienced evaluator via B-mode ultrasound imaging using a 7.5 MHz linear probe (MySono U6, Samsung, Sao Paulo, Brazil). The mCSA acquisition and quantification technique validated by Reeves et al. was used (Reeves *et al*., 2004).

Participants were instructed to refrain from vigorous physical activity for at least 72 hours before each assessment. At arrival, participants laid in a supine position, and a point was measured at the midpoint of the greater trochanter and the lateral epicondyle, representing 50% femur length. This was used as a reference for image acquisition. From there, successive markings were made every 2 cm in the medial and lateral directions to guide the displacement of the ultrasound probe. This occurred during a 15-minute period in which participants laid still to allow for tissue fluid shift stabilization. Ultrasound images were captured sequentially using the aforementioned markings as a guide for displacement of the probe in the sagittal plane. Imaging began at the medial-most mark, located over the rectus femoris muscle. Water-soluble transmission gel was applied to ensure acoustic coupling of the probe without skin compression. Obtained images were manually rotated and sequentially overlaid to create a panoramic image of the CSA of the entire muscle. The ImageJ “polygonal” tool was used to circle and calculate VL mCSA, while excluding bone and connective tissue. The typical error between two image acquisitions and quantifications separated by 72 hours was 0.52 cm^2^ (2.47%).

### Muscle tissue processing and analytical techniques

#### Vastus lateralis muscle biopsies

Muscle tissue samples were obtained from the VL via percutaneous muscle biopsies performed using the Bergström method. The area around the sampling site was cleaned with an antiseptic wash and injected with 2-3 mL of 1% xylocaine (lidocaine) for local anesthesia. A small incision was then made with a surgical scalpel and inserted the biopsy needle (5-gauge Bergström needle with manual suction) to a depth of ∼4 cm to remove ∼100 mg of muscle tissue. The incision was then closed and covered with bandages while the sampled tissue was cleaned of blood and teased of connective tissue. The tissue used for ribosomal analysis herein (∼30 mg) was allocated to cryotubes and frozen in liquid nitrogen for analysis. Tissue utilized for the analysis of fCSA was positioned in optimal cutting temperature (OCT) solution. These samples were oriented such that fibers were perpendicular to the horizontal surface then frozen in liquid nitrogen-cooled isopentane for immunohistochemical analysis. All samples were then stored at -80° C until later processing.

#### Ribosome enrichment process

Figure 1 depicts a workflow of included participants, ribosome enrichment process, and analytical methods utilized for this study. The following protocol is a modification of the ribosome isolation protocol from Godwin et al. (Godwin *et al*., 2025), which was used to sufficiently isolate ribosomes from C2C12 myotubes. The base buffer used for these experiments (Ribo buffer) consisted of 50 mM Tris-HCl, 250 mM KCl, and 25 mM MgCl_2_ in diethyl pyrocarbonate (DEPC) treated water. The buffer used for homogenization (Ribo homogenization buffer) consisted of 0.25 mM dithiothreitol (DTT), 1 mg/mL cycloheximide, 80 U/mL RNase inhibitor (RiboLock; cat no. EO0382; Thermo Fisher Scientific; Waltham, MA, USA), 0.5% Triton-X. Following tissue collection, ∼30 mg of tissue was weighed on an analytical scale. This tissue was then homogenized in Ribo buffer (10 μL/mg tissue) with tight-fitting plastic pestles in 1.7 mL tubes. Lysates were then centrifuged at 12,000 *g* for 10 mins at 2° C to pellet insoluble protein fraction, cell debris, and nuclei. Supernatants (soluble cell fraction) were then overlaid onto a 20% sucrose gradient (in Ribo buffer) in 13.2 mL polyallomer tubes (Seton Scientific; cat no. 5030). Tubes were then placed in a swinging bucket TH-641 rotor (Thermo Fisher; cat no. 54295) and centrifuged at 100,000 *g* (28,200 RPM) for 3 hours at 2°C in an ultracentrifuge (Thermo Fisher; cat no. 75000080).

**Figure 1.**
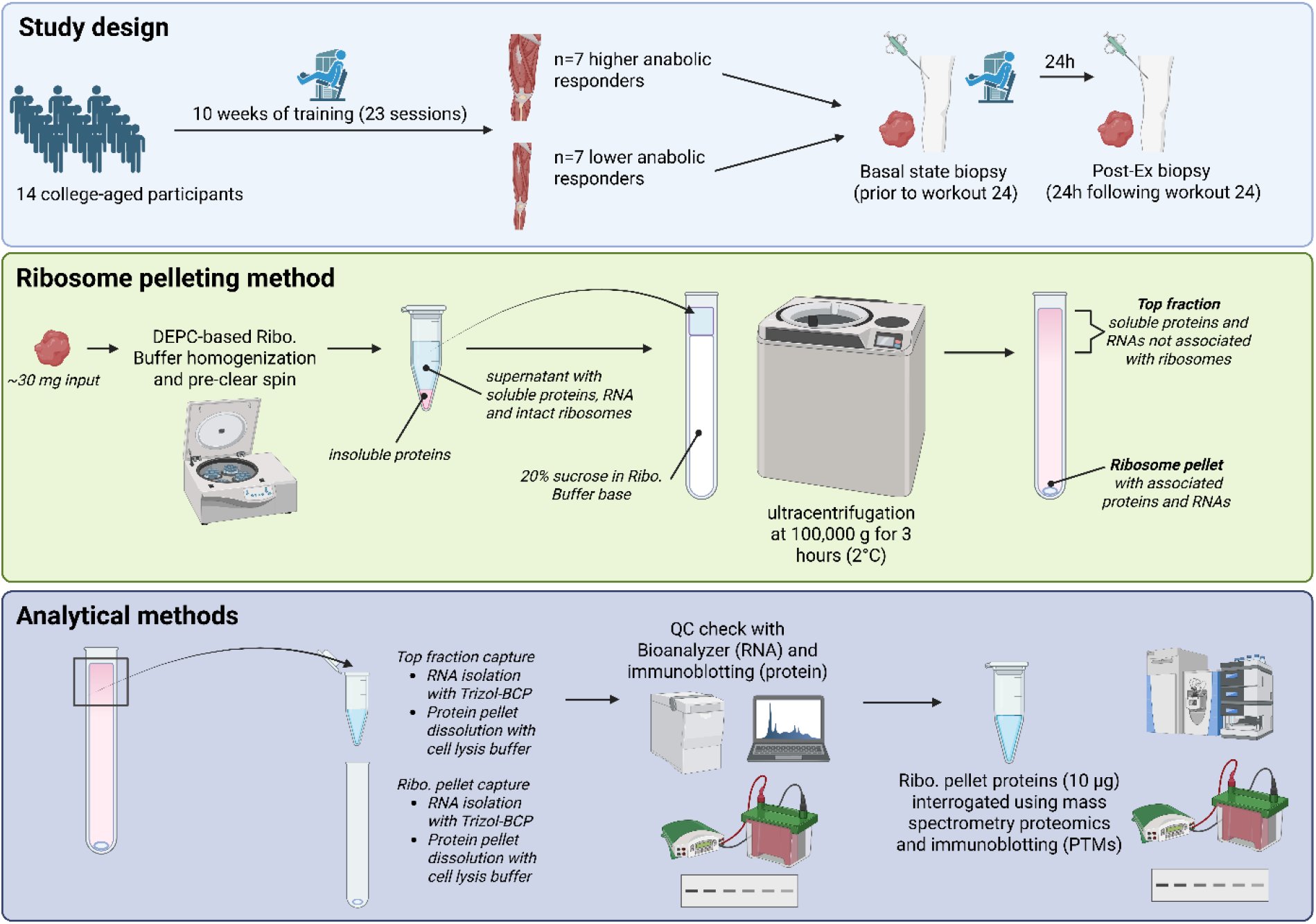
Study design. Legend: Summary of study design. Schematic (drawn using Biorender.com) illustrates study logistics and participant number for this study. More details regarding study design can be found in text. Abbreviations: DEPC: diethyl pyrocarbonate; Ribo: ribosome; BCP: 1-bromo-3-chloropropane; QC: quality control; PTM: post-translational modifications.

Following ultracentrifugation, the top 1 mL of the sucrose solution was collected and placed in a 1.7 mL tube with 500 μL of Trizol (ThermoFisher; cat no. 15596026) and the remainder of the sucrose solution was decanted as waste. 500 μL of Trizol was then triturated in the bottom of the polyallomer tube to dislodge the ribosome pellet. This ribosome pellet-containing Trizol was then placed in a 1.7 mL tube. These tubes were then frozen at -20°C overnight.

#### Immunohistochemistry for the fiber cross-sectional area

OCT-preserved samples were sectioned at a thickness of 14 μm using a cryostat (Leica Biosystems, Buffalo Grove, IL, USA) and adhered to positively charged slides. These prepared slides were held at -80°C until being batch-processed for immunohistochemical analysis for the quantification of cross-sectional area. To begin batch-processed staining, slides were elevated and air dried for ≥2 hours at room temperature. Once dry, sections were then fixed in acetone at -20°C for 5 minutes. Endogenous peroxidases were then blocked with 3% H_2_O_2_ for 10 min at room temperature. Slides were then incubated for 1 minute with an autofluorescence quenching agent (TrueBlack, cat no. 23007, Biotium, Fremont, CA, USA). Slides were subsequently blocked for 1 hour with a 5% goat serum, 2.5% horse serum, 0.1% Triton-X solution in PBS at room temperature. After blocking, slides were incubated overnight at 4°C with a 1:1000 rabbit anti-dystrophin primary antibody (cat no. GTX57970; GeneTex, Irvine, CA, USA) in 2.5% horse serum solution in PBS. The following day, sections were washed and then incubated for 1h in a 1:250 goat anti-rabbit IgG DyLight488 (cat no. DI-1488; Vector Laboratories, Newark, CA, USA) in PBS. Slides were then stained with DAPI (4’,6-diamidino-2-phenylindole; cat no. D3571; Thermo Fisher Scientific) for 10 min at room temperature. Slides were then mounted with glass coverslips using a 1:1 PBS:glycerol mounting solution. Sections were then stored in the dark at 4°C until imaging was conducted. Digital images were captured with a fluorescence microscope at x20 magnification (Nikon Instruments; Melville, NY, USA). All areas selected for analyses were free of freeze-fracture artifacts, and at least 50 fibers per sample were quantified (Mackey *et al*., 2009).

#### RNA isolation

Following an overnight hold at -20°C, 100 μL of 1-bromo-3-chloropropane (BCP) was added to each tube after which the tubes were shaken and allowed to sit at room temperature for 3 minutes. Tubes were then spun at 12,000 *g* for 10 minutes at 2°C. The top, aqueous, fraction was then extracted for use in RNA isolation, and the remainder of this sample was used for protein isolation. Notably, given the robust difference in starting volumes between the top fraction and the ribosome pellet (1.5 mL versus 0.5 mL respectively), 1 mL of the aqueous phase was used for RNA isolation from the top fraction whereas 300 μL was used for RNA isolation in the ribosome pellet fraction. 250 μL of isopropyl alcohol and 0.5 μL of glycogen were then added to each sample, whereafter samples were once again shaken and incubated on ice for 10 minutes. Samples were then centrifuged at 12,000 *g* at 2°C for 10 minutes. An RNA pellet was generated from this spin, and the supernatant was decanted as waste. Two consecutive washes were then conducted where 500 μL of 75% ethanol in DEPC water was added to each tube, which were then centrifuged at 7,500 *g* at 2°C for 5 minutes.

Supernatants were discarded as waste, and tubes were inverted for 10 minutes to allow samples to dry. Finally, RNA pellets were reconstituted in 30 μL of DEPC water. RNA samples were then stored at -80°C until further analysis.

#### Quantification of total RNA, 18S rRNA, and 28S rRNA

RNA concentration for each sample was determined using a NanoDrop Lite (ThermoFisher; cat no. NDLPLUSGL) with reads performed in duplicate. 18S and 28S rRNA concentrations were examined using a microfluidic gel electrophoresis (MFGE) via a commercially available kit (Agilent; Santa Clara, CA, USA; cat no. 5067-1511) measured with the Agilent 2100 Bioanalyzer system. This involved first decontaminating the machine’s electrodes using RNase Away (Thermo Fisher Scientific; cat no. 7003) followed by DEPC water. The RNA Nano gel matrix was then filtered and mixed with the provided dye concentrate according to manufacturer’s specifications. The chip was prepared by loading 9 μL of gel-dye mix into the designated wells, followed by 5 μL of RNA marker and 1 μL of either RNA ladder or sample in their respective wells. The chip was vortexed at 2400 RPM for 1 minute and analyzed within 5 minutes.

#### Protein isolation

Following BCP phase separation, the remaining phases following the removal of the aqueous phase were used for protein isolation. DNA and protein were first separated by adding 300 μL of ethanol to each sample, shaking, incubating at room temperature for 3 minutes, and centrifugation at 5,000 *g* for 10 minutes at room temperature. The resultant sample consisted of a DNA pellet and a Trizol/protein containing supernatant. This supernatant was then removed and placed into new 2 mL tubes. 650 μL of ethanol, 100 μL of BCP, and 600 μL of ethanol, were then added to the samples, where each step was followed by vigorous vortexing. Samples were then centrifuged at 12,000 *g* for 5 minutes at room temperature. The aqueous phase was then decanted as waste, and another 700 μL of ethanol was added to each sample followed by vortexing. Samples were once again centrifuged at 12,000 *g* for 5 minutes at room temperature, generating a protein pellet. Tubes were then inverted for 5 minutes to allow the newly formed protein pellet to dry. Samples were then resuspended in general lysis buffer (Cell Signaling; Danvers, MA, USA; cat no. 9803). Due to variations in the size of the generated protein pellets, 1 mL of general cell lysis buffer was used to suspend the protein pellet generated from the top fraction, while 100 μL was used to suspend the protein pellet generated from the ribosome pellet fraction. Tight fitting plastic pestles were used to homogenize protein pellets where needed. Samples were then vortexed and held at -80°C until further analysis.

#### Western blotting

Subsequent protein lysates were processed for total protein using a commercially available BCA protein assay kit (Thermo Fisher Scientific, Waltham, MA, USA; Cat. No. A55864) and spectrophotometer (Agilent Biotek Synergy H1 hybrid reader; Agilent, Santa Clara, CA, USA). Lysates were prepared for western blotting using 4x Laemmli buffer and deionized water (diH_2_O) at equal protein concentrations (0.25 μg/μL). 15 μL of samples were pipetted onto SDS gels (4-15% Criterion TGX Stain-free gels, Bio-Rad Laboratories; Hercules, CA, USA), and proteins were separated by electrophoresis at 200 V for 45-50 min. Proteins were then transferred to methanol-preactivated PVDF membranes (Bio-Rad Laboratories) for 2 hours at 200 mA, Ponceau stained for 10 minutes, washed with diH_2_O for ∼30 seconds, dried, and digitally imaged (ChemiDoc Touch, Bio-Rad). Following Ponceau imaging, membranes were reactivated in methanol, blocked with non-fat bovine milk for ≥1 hour, and washed 3×5 minutes in Tris-buffered saline with tween 20 (TBST). Membranes were incubated with primary antibodies (1:1000 v/v dilution in TBST with 5% (w/v) bovine serum albumin (BSA)) on a rocker overnight at 4°C. Antibodies against GAPDH (GeneTex; Irvine, CA, USA; cat no.

GTX100118), and a cocktail containing RPS3 (Cell Signaling Technologies; cat no. 9538S), RPL5 (Cell Signaling Technologies; cat no. 14568S), and RPL11 (Cell Signaling Technologies; cat no. 18163S) were used to detect these protein abundances. Following primary antibody incubations, membranes were washed 3×5 minutes in TBST and incubated for 1 hour with HRP-conjugated anti-rabbit IgG (Cell Signaling Technology; cat No. 7074) at a 1:2000 dilution in TBS-T with 5% BSA (w/v). Membranes received a final set of 3×5-minute washes in TBST, then developed using chemiluminescent substrate (Millipore, Burlington, MA, USA), and then digitally imaged. Notably, formal densitometry analysis and subsequent statistical testing were not performed on western blotting images as the generated images were used only to confirm enrichment of ribosomal proteins in the ribosome pellet fraction and enrichment of GAPDH in the top fraction.

#### Peptide isolation and proteomics

Each sample (10 µg) was prepared for LC-MS/MS analysis using EasyPep Mini MS Sample Prep Kit (Thermo Fisher Scientific; cat no. A4006). In brief, samples were transferred into new microcentrifuge tubes and final volumes were adjusted to 100 µL with general cell lysis buffer (Cell Signaling). Reduction solution (50 µL) and alkylation solution (50 µL) were added to samples, gently mixed, and incubated at 95°C using a heat block for 10 minutes to reduce and alkylate samples. After incubation, the sample was removed from the heat block and cooled to room temperature. A reconstituted enzyme Trypsin/Lys-C Protease solution (25 µL) was added to the reduced and alkylated protein sample and incubated with shaking at 37°C for 2 hours to digest proteins. Digestion stop solution (25 µL) was then added to samples and peptides were cleaned using a peptide clean-up column according to the kit instructions. An externally calibrated Thermo Orbitrap Exploris 240 (high-resolution electrospray tandem mass spectrometer) was used in conjunction with Dionex UltiMate3000 RSLC Nano System (Thermo Fisher Scientific). Samples were aspirated into a 20 µL loop and loaded onto the trap column (Thermo Acclaim PepMap 100 nanoviper tubing 75 µm i.d. × 2 cm). The flow rate was set to 300 nL/min for separation on the analytical column (Easy-Spray pepmap RSLC C18 50 µM × 15 cm Thermo Fisher Scientific). Mobile phase A was composed of 99.9% H_2_O (EMD Omni Solvent; Millipore, Austin, TX, USA), and 0.1% formic acid and mobile phase B was composed of 80% acetonitrile and 0.1% formic acid. A 135-minute linear gradient from 3% to 50% B was performed. The eluent was directly nanosprayed into the mass-spectrometer. During chromatographic separation, the Orbitrap Exploris was operated in a data-independent mode and under direct control of the Thermo Excalibur 4.4.16.14 (Thermo Fisher Scientific). The MS data were acquired from (380 to 985 m/z) at 60,000 resolution. Data independent acquisition MS2 were acquired with an isolation window of 10 m/z and an overlap of 1 m/z with collision energy fixed at 28 and 15,000 resolution. All measurements were performed at room temperature, three technical replicates were run for each sample, and mean values were used for obtaining expression values relative to total spectra.

#### DIA-NN data processing and protein annotation

Raw.dia files acquired from mass spectrometry were processed using DIA-NN (v 2.2.0) (Demichev *et al*., 2020). All raw files were analyzed jointly in a single batch to enable retention-time alignment and match-between-runs (MBR) quantification. In silico digestion was performed with trypsin-P specificity, allowing one missed cleavage. Carbamidomethylation of cysteine was specified as a fixed modification, while oxidation of methionine was included as a variable modification, with one variable modification permitted per peptide. N-terminal methionine excision was enabled. Peptide identification was guided by the DPHLv2 spectral library (Xue *et al*., 2023). For protein reannotation, a reviewed *Homo sapiens* UniProt reference proteome (release 2020-04) supplemented with indexed retention-time (iRT) calibration peptides, reversed decoys, and common contaminants were used.

### Statistical analyses

*RNA and VL size changes*. All data except for proteomic data were checked for normality via Shapiro-Wilk test prior to statistical analysis. After confirming that data were normally distributed, total RNA concentration was analyzed via a paired samples t-test, while VL mCSA and fCSA change scores were compared via independent samples t-tests. VL mCSA and fCSA, along with 18S and 28S rRNA abundance changes were assessed via two-way ANOVA (training responsiveness*time). Significance level was set to P<0.05 for all comparisons listed above.

#### Bioinformatics for proteomic profiling of ribosomal proteins

Relative protein expression values were obtained from DIA-NN precursor intensities, which were summed to protein level, normalized to the total summed spectra, and log_2_-transformed. Proteins missing in ≥50% of samples were excluded (Shi *et al*., 2021; Shen *et al*., 2024), and remaining missing values were imputed with zeroes. Linear modeling was performed using limma with duplicateCorrelation to account for repeated measures. Group, time, and interaction contrasts were evaluated; significance was set at P<0.01. Proteins were annotated using UniProt.ws and cross-referenced with a custom mapping file.

## RESULTS

### Ribosomal proteins and rRNAs are only present in the ribosome pellet fraction

Preliminary validation experiments indicated that RNA concentrations were higher in the ribosome pellet fraction (139.1 ng/μL) versus the top fraction (71.5 ng/μL) of the sucrose solution (P<0.001; Figure 2A). Furthermore, when MFGE was performed on each fraction 5.8S, 18S, and 28S rRNA peaks were only detected in the ribosome pellet fraction (Figure 2B).

**Figure 2.**
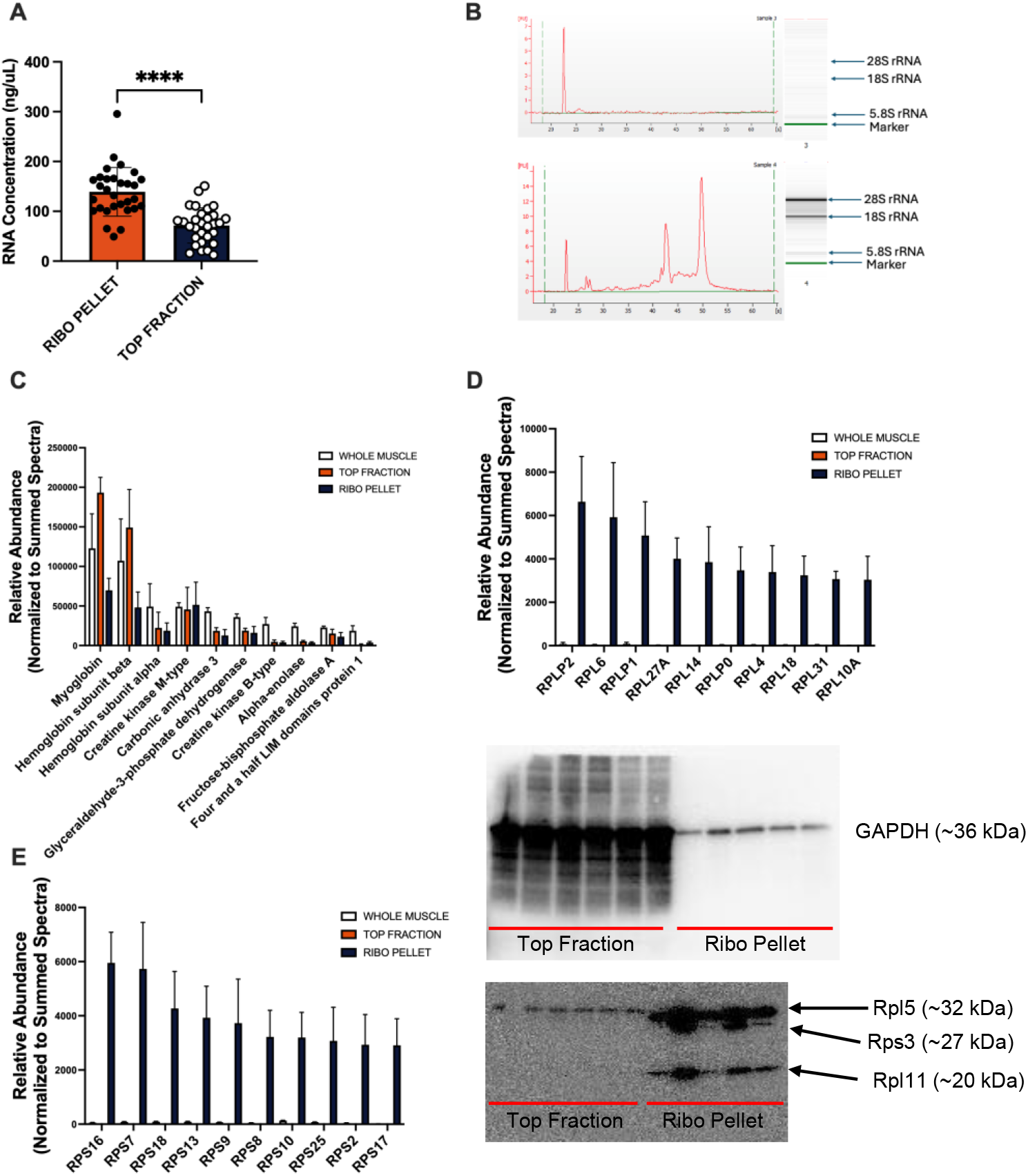
Validation of ribosome enrichment protocol. Legend: Validation of Ribosome pellet fidelity. Panel A: RNA concentration assessment via spectrophotometry. Panel B: Assessment of rRNA content via fluorometry-based microfluidic chip electrophoresis in a single participant. Panel C: Protein abundances of top 10 detected proteins by abundance in whole muscle compared to the top fraction isolate and ribosome pellet isolate in 3 participants not involved in responder analysis. Panel D: Top large subunit ribosomal proteins (RPL) detected in the ribosome pellet as compared to whole muscle and the top fraction isolate in 3 participants not involved in responder analysis. Panel E: Top small subunit ribosomal proteins (RPS) detected in the ribosome pellet as compared to whole muscle and the top fraction isolate in 3 participants not involved in responder analysis. Panel F: Western blot image comparing GAPDH expression in the top fraction isolate versus the ribosome pellet. Panel G: Western blot image comparing a ribosomal protein cocktail (RPL5, RPS3, and RPL11) in the top fraction isolate versus the ribosome pellet.

Interestingly, despite suitable RNA concentrations in the top and pellet fractions, qPCR was performed, the genes *MYH2, TTN*, and *GAPDH* were expressed only in the ribosome pellet fraction and a whole muscle comparator (data not shown), likely indicating some mRNA contamination of the ribosome pellet fraction. Notwithstanding, further evidence of effective separation of ribosomal constituents to the ribosome pellet fraction comes from protein interrogations. When proteomics was performed on samples from ribosome pellet fractions, top fractions, and whole muscle lysate comparators, non-ribosomal proteins were generally more abundant in the whole muscle and top fraction samples than the ribosome pellet fractions (Figure 2C). Furthermore, it was observed that the top 10 most abundant small ribosomal proteins, and the top 10 most abundant large ribosomal proteins were almost exclusively present in the ribosome fraction (Figure 2 D-E). To follow up on this finding, western blotting was performed on GAPDH and a cocktail of three ribosomal proteins (RPS3, RPL5, RPL11). Similar to proteomic data, it was observed that GAPDH was highly enriched in the top fraction and not the ribosome pellet fraction, whereas the ribosomal protein cocktail signal was highly enriched in the ribosome pellet fraction and not the top fraction (Figure 2 F-G). Raw abundances of all proteins identified from the whole-muscle samples, top fractions, and ribosome pellets are presented in the Supplemental data found at synapse.org; DOI: https://doi.org/10.7303/syn70283453.

### Higher versus lower responder characteristics

When analyzing data from the 14 participants used for higher and lower responder analysis, VL mCSA demonstrated main effects of time and training response (P≤0.001) along with a significant training response*time interaction (P<0.001) where higher responders significantly increased this metric while lower responders did not (Figure 3A). Similarly, VL fCSA demonstrated a main effect of time (P=0.046) and a significant training response*time interaction (P=0.044) with higher responders once again increasing in this metric while lower responders did not (Figure 3B). Furthermore, independent samples t-tests revealed that high responders had significantly higher change scores of VL mCSA (P<0.001; Figure 3C) and VL fCSA (P<0.05; Figure 3D).

**Figure 3.**
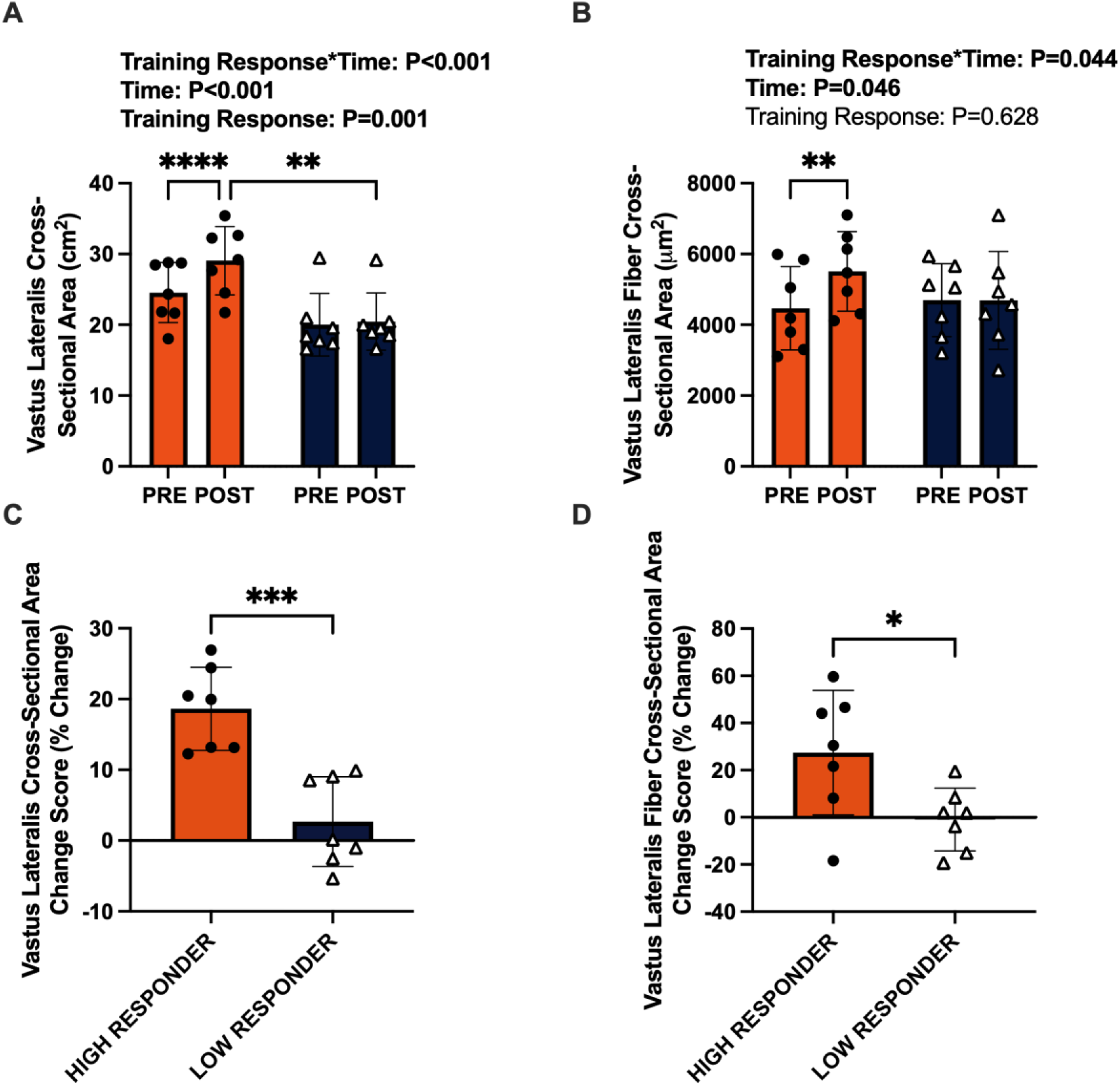
Characteristics of higher versus lower responders to resistance training. Legend: Muscle size characteristics of higher (n=7) versus lower (n=7) hypertrophic responders to 10 weeks of resistance training. Panel A: Ultrasound-derived vastus lateralis muscle cross-sectional area before (PRE) and after 10 weeks of RT (POST). Panel B: Vastus lateralis fiber cross-sectional area before (PRE) and after 10 weeks of RT (POST). Panel C: Change score in ultrasound-derived vastus lateralis muscle cross-sectional area from before (PRE) to after 10 weeks of RT (POST). Panel D: Change score in vastus lateralis fiber cross-sectional area from before (PRE) to after 10 weeks of RT (POST). Symbols: *, P<0.05; **, P<0.01, ***, P<0.005; ****, P<0.0001.

### Individual ribosomal proteins largely do not differ between responder cohorts

When analyzing data from the 14 participants used for higher and lower responder analysis, abundances of the 18S and 28S rRNA subunits did not show main effects of time (P≥0.648), training response cohort (P≥0.650), or a training response*time interaction (P>0.752) in response to a single bout of RT (Figure 4A-B). Proteomics on enriched ribosomes from these participants indicated that no large or small ribosomal proteins showed main effects of time (P≥0.034), training response (P>0.028), or a training response*time interaction (P>0.094; The top 10 large and small ribosomal proteins by total abundance from these analyses are shown in Figure 4C-D, and the remaining ribosomal protein data as well as other supplemental data associated with these analyses can be found at synapse.org; DOI: https://doi.org/10.7303/syn70283453).

**Figure 4.**
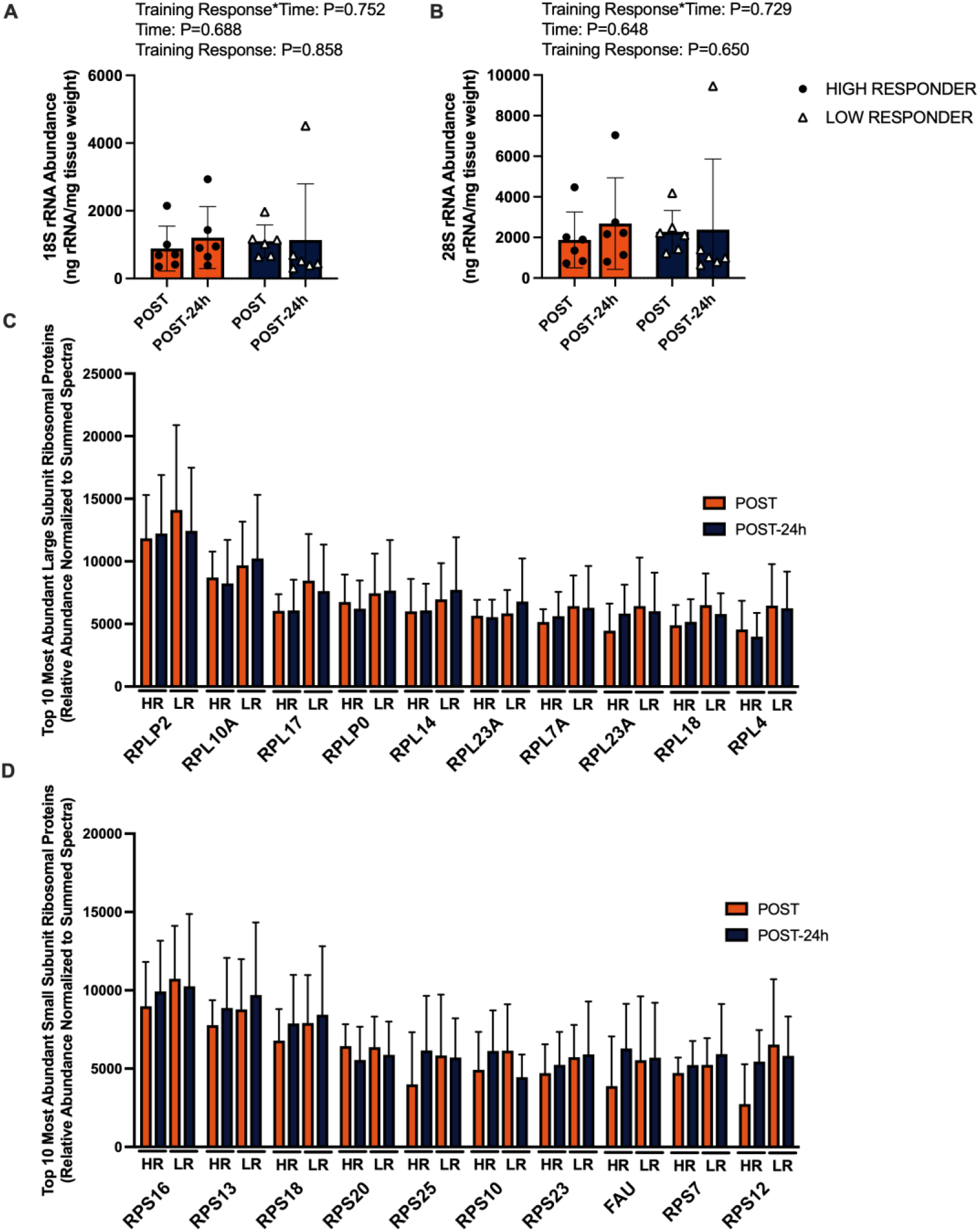
Ribosomal characteristics of higher versus lower responders. Legend: Ribosome characteristics of higher (HR, n=7) versus lower (LR, n=7) hypertrophic responders to 10 weeks of resistance training prior to (POST) and following RT session 24 (POST-24h). Panel A: 18S rRNA subunit abundance. Panel B: 28S rRNA subunit abundance. Panel C: Top 10 large subunit proteins (RPL) by abundance in the ribosome pellet. Panel D: Top 10 small subunit proteins (RPS) by abundance in the ribosome pellet.

## DISCUSSION

In agreement our hypothesis, the utilized homogenization and ultracentrifugation techniques enable the enrichment of ribosomes from a relatively low amount of human skeletal muscle tissue. However, contrary to our hypothesis, the responder analysis suggests that RT hypertrophic response heterogeneity does not coincide with differential ribosomal protein profile shifts; this latter finding is likely being in part due to sample size and biopsy sampling limitations which will be discussed in the following paragraphs.

This methodological approach indicates that ribosomes can be successfully enriched from a limited amount of human muscle biopsy sample (∼30 mg), and expands upon our recent publication by Godwin et al. (Godwin *et al*., 2025) who effectively isolated ribosomes from C2C12 myotubes. Notably, the present protocol differs from our past publication in that the sucrose gradient was reduced from a 30% to 20% solution given that the former proved to be too dense to pellet ribosomes from tissue samples, as evidenced by the absence of rRNA peaks following ultracentrifugation (data not shown). However, certain caveats to our approach should be noted. First, unlike polysome fractionation, which has long been performed in rodent skeletal muscle/cell cultures (Srivastava, 1969; Roberts *et al*., 2012) and human primary muscle cells (Mei *et al*., 2022), the current approach indiscriminately pellets all ribosomes into a single fraction. Although the fidelity of this method appears high (Figure 2), the ribosome pellet did contain select mRNAs and non-ribosomal proteins. Indeed, it is possible that these targets may either represent mRNAs actively undergoing translation or nascent proteins being synthesized, albeit this remains speculative as profiling of mRNAs directly associated with ribosomes (Ribo-Seq) would be required to confirm translational activity (Li *et al*., 2017; Mahmassani *et al*., 2021). Nevertheless, this protocol provides a unique, relatively low-cost approach to enrich and study ribosomal proteins from limited human muscle tissue and can serve as guidance for further optimization of sucrose gradient and ultracentrifugation parameters (speed and duration) for researchers interested in fine-tuning the technique.

Proteomic interrogation of the ribosome pellet identified 74 distinct ribosomal proteins and four analogous variants (RPL36AL, RPS27L, RPL3L, RPLP0P6), representing approximately 92% coverage of the 80 putative ribosomal proteins (Anger *et al*., 2013; Gupta & Warner, 2014; de la Cruz *et al*., 2015; Khatter *et al*., 2015). Specifically, 31 ribosomal proteins from the 40S subunit (RPS) and 43 from the 60S subunit (RPL) were quantified. In comparison, whole-muscle lysate proteomics detected 71 total cytosolic ribosomal proteins (30 RPS, 41 RPL), indicating that this enrichment method improved the detection of three ribosomal proteins. Perhaps more notable, however, is that the pelleting and enrichment methods increased the relative enrichment of RPLs and RPSs by ∼164-fold and ∼71-fold respectively when compared to conventional homogenization (Top 10 RPLs and RPSs depicted in Figure 2D and 2E). Thus, this further supports that our methodological approach enables future avenues of ribosome profiling from limited tissue inputs.

In relation to our RT response heterogeneity analysis, ribosomal protein composition and surrogate markers of ribosomal content (i.e., rRNAs) did not differ prior to or 24 hours following a single RT bout (session 24) between the higher and lower hypertrophic responders who had completed 23 prior RT sessions over a 10-week period. Although this was somewhat unexpected given prior associations between ribosome content and RT responsiveness (Stec *et al*., 2016; Hammarstrom *et al*., 2020; Hammarstrom *et al*., 2022), the number of participants as well as the analysis of only pre- and post-session 24 are significant limitations. Pe-intervention muscle biopsy samples would have provided tremendous insight, albeit tissue was used in prior publications and were not available for analysis herein. Indeed, others have reported that ribosome content alterations within the first 1-2 weeks of RT demonstrate high positive correlations to indices of skeletal muscle hypertrophy (Hammarstrom *et al*., 2022). Hence, while not assessed, it is indeed possible that RP alterations may have occurred over the 10-week training period but that these adaptations may have been masked due to high levels of ribosome turnover that occur quickly in response to mechanical overload (Brook *et al*., 2017; Hammarstrom *et al*., 2022). Furthermore, Nader et al. (Nader *et al*., 2014) reported that 12 weeks of RT alters the expression pattern of genes associated with exercise adaptation as compared to a single naïve RT bout. This suggests that even if RP variation is a factor in response heterogeneity in the early stages of RT and that adopting a trained molecular phenotype might alter RP production such that these changes are undetectable following 10 weeks of RT. Additionally, our proteomic approach assessed only overall protein abundance of each RP, but did not account for post-translational modifications that may influence ribosome function and specialization (Liew & Gornall, 1973; Shin *et al*., 2009; Shirai *et al*., 2010; Matsuo *et al*., 2017; An & Harper, 2020; Montellese *et al*., 2020; Pletnev *et al*., 2022; Ni *et al*., 2023). Thus, future investigations with more sampling time points implementing our ribosome enrichment technique will enable the investigation of these potential avenues.

## Conclusions

This study establishes a novel, cost-effective method to enrich ribosomes from human skeletal muscle using a relatively small tissue input (∼30 mg). Our method enabled a robust enrichment of ribosomal proteins compared to conventional tissue homogenization. Our preliminary analysis suggests individual ribosomal protein alterations do not coincide with RT hypertrophic response heterogeneity. However, given the sample size and biopsy sampling limitations discussed herein, we cannot exclude the possibility that such differences influence the adaptive response to resistance training and future studies implementing similar analytical approaches are needed to provide further clarity.

## ADDITIONAL INFORMATION

## Acknowledgments

Supplemental data files can be found online at synapse.com (DOI: https://doi.org/10.7303/syn70283453). Other raw data related to the current study outcomes will be provided upon reasonable request by emailing the corresponding author.

## Clinical Trial Registration

The human subjects portion of this study was registered as a clinical trial in the Brazilian Registry of Clinical Trials (RBR – 57v9mrb).

## Ethics Approval Statement

Study protocols were carried out in accordance with the most recent version of the declaration of Helsinki. All study procedures were approved by the Ethics Committee of the Federal University of São Carlos (no. 56259622.0.0000.5504).

## Patient Consent Statement

All participants in this study provided verbal and written consent in accordance with the above IRB approval.

## Author Contributions

JMM and MDR primarily drafted the manuscript; JMM and MDR prepared figures; JMM, SCN, and CBM primarily carried out laboratory based assays; MDB performed mass spectrometry for the measurement of the ribosomal proteome; CGU performed DIA-NN data processing and annotation; CAL provided funding, oversight, and gathered approval for the human subjects portion of this study; JSG and GAN provided intellectual feedback for the duration of the project. All co-authors listed assisted with revising and editing the manuscript, and all co-authors approve the final version.

